# High-throughput gene expression analysis with TempO-LINC sensitively resolves complex brain, lung and kidney heterogeneity at single-cell resolution

**DOI:** 10.1101/2024.08.03.606484

**Authors:** Dennis J. Eastburn, Kevin S. White, Nathan D. Jayne, Salvatore Camiolo, Gioele Montis, Seungeun Ha, Kendall G. Watson, Joanne M. Yeakley, Joel McComb, Bruce Seligmann

## Abstract

We report the development and performance of a novel genomics platform, TempO-LINC, for conducting high-throughput transcriptomic analysis on single cells and nuclei. TempO-LINC works by adding cell-identifying molecular barcodes onto highly selective and high-sensitivity gene expression probes within fixed cells, without having to first generate cDNA. Using an instrument-free combinatorial-indexing approach, all probes within the same fixed cell receive an identical barcode, enabling the reconstruction of single-cell gene expression profiles across as few as several hundred cells and up to 100,000+ cells per run. The TempO-LINC approach is easily scalable based on the number of barcodes and rounds of barcoding performed; however, for the experiments reported in this study, the assay utilized over 5.3 million unique barcodes. TempO-LINC has a robust protocol for fixing and banking cells and displays high-sensitivity gene detection from multiple diverse sample types. We show that TempO-LINC has an observed multiplet rate of less than 1.1% and a cell capture rate of ∼50%. Although the assay can accurately profile the whole transcriptome (19,683 human or 21,400 mouse genes), it can be targeted to measure only actionable/informative genes and molecular pathways of interest – thereby reducing sequencing requirements. In this study, we applied TempO-LINC to profile the transcriptomes of 89,722 cells across multiple sample types, including nuclei from mouse lung, kidney and brain tissues. The data demonstrated the ability to identify and annotate at least 50 unique cell populations and positively correlate expression of cell type-specific molecular markers within them. TempO-LINC is a robust new single-cell technology that is ideal for large-scale applications/studies across thousands of samples with high data quality.

## Introduction

Single cell gene expression profiling assays are critical tools for identifying functional subtypes of cells, as well as changes within cells resulting from pathology or treatment (Carangelo et al., 2022; Tabula Sapiens Consortium et al., 2022; Wen et al., 2022). Current methods have limitations in one or more key areas: cost, low sample throughput, poor sample recovery, providing cell-associated reads that either fail to map to transcripts or map to uninformative genes, reliance on reverse transcription and/or insufficient sensitivity to consistently measure lowly expressed genes from single cells - preventing measurements of many key biomarkers and molecular pathways (Conte et al., 2024; Ding et al., 2020; Yamawaki et al., 2021). The above limitations have contributed to most single-cell sequencing studies being restricted to smaller numbers of samples and have slowed their wider scale application in biopharma, clinical, translational and applied markets where large numbers of samples become cost prohibitive (Van de Sande et al., 2023).

Combinatorial indexing or split pool barcoding methods have been introduced to address some issues related to the high cost and limitations on scalability associated with single-cell transcriptomics (Cao et al., 2017, 2019; Rosenberg et al., 2018; Srivatsan et al., 2020). With the current combinatorial single-cell approaches, reverse transcription and barcoding are performed directly on nucleic acids within fixed cells. Following each round of barcode labeling, cells are pooled together and then randomly redistributed into unique wells containing one of the next round’s barcodes. Multiple rounds of barcoding ensure that each cell receives a unique barcode combination, thereby enabling transcriptome profiles to be mapped to individual cells following sequencing and demultiplexing. Plate-based split pool barcoding methods have highly scalable single-cell throughput that is predominantly limited by the number of unique barcodes used in each round of barcoding and the number of rounds of barcoding. The use of the split pool workflow first-round barcode as a unique sample identifier also enables simplified labeling and multiplexing of greater numbers of samples. The aforementioned advantages, combined with the instrument-free nature of the workflows serve to make these methods generally less expensive than droplet-based or microwell approaches (De Simone et al., 2024). However, due to the reliance on reverse transcription, simultaneously optimizing fixation for both robust cell integrity and high-sensitivity cDNA synthesis can be challenging with combinatorial split pool methods. Compatibility with formalin-fixed, paraffin-embedded (FFPE) isolated samples has also not been demonstrated with these approaches (De Simone et al., 2024).

Alternative methods for bulk sample expression profiling that do not rely on the reverse transcription step of traditional RNA-Seq have been developed. One such method, TempO-Seq, is based on detector oligonucleotide (DO) probes that specifically hybridize to adjacent regions within mRNAs (Cannizzo et al., 2022; Trejo et al., 2019; Yeakley et al., 2017). When properly hybridized to the complementary regions within mRNAs, the 3’ hydroxyl group on the Downstream Detector Oligo (DDO) is ligated to the 5’ phosphate of the Upstream Detector Oligo (UDO) thereby forming an amplifiable ligation product. Subsequent PCR amplification is performed using universal primer landing sites common to all DDOs and UDOs. During this amplification, sample indices and sequencing adaptors are also added, enabling short read sequencing to identify and count the numbers of each probe within a library. This approach has a number of key advantages including but not limited to: 1) lack of 3’ bias, allowing measurement of any sequence where there may be a gene fusion or mutation, including expressed single base variants and alternatively spliced isoforms, 2) works on poor quality RNA, fixed and FFPE samples and H&E stained FFPE samples (Cannizzo et al., 2022; Trejo et al., 2019), 3) simple bioinformatic mapping, 4) content flexibility from custom gene panels up to the entire transcriptome, 5) the ability to quantitatively attenuate highly expressed genes, thus increase effective dynamic range and sensitivity and/or the number of samples that can be multiplexed into a single sequencing run (Yeakley et al., 2017), and 6) low per sample reagent costs.

Here, we report on the development and performance of TempO-LINC, a novel genomics platform for single-cell gene expression profiling derived from and using the same commercial and custom assay content as TempO-Seq. TempO-LINC combines the benefits of the TempO-Seq assay, including the lack of a reverse transcription step, with a combinatorial split pool barcoding approach to provide transcriptome-wide gene expression measurements at single-cell resolution. TempO-LINC has a user friendly and instrument-free workflow that is highly scalable across cells and samples. Significantly, we show robust assay performance and gene detection rates across a variety of cells and sample types that enabled the accurate analysis of complex tissue heterogeneity.

## Results

### Transcriptional Profiling with the TempO-LINC Workflow

To enable high-throughput single-cell resolution with the TempO-Seq gene expression assay, we sought to develop a combinatorial split pool-based approach to add cell-identifying molecular barcodes to the 5’ end of ligated DO probe pairs (Figure 1A). To accomplish this, we first bulk phosphorylated the DDOs contained in the DO sets of commercial TempO-Seq assays, then implemented and optimized an *in situ* hybridization version of TempO-Seq developed for use with formaldehyde fixed cells (Trejo et al., 2019). After formaldehyde fixation, cells are permeabilized and incubated with the fully phosphorylated DOs targeting the transcriptome. Following overnight hybridization to mRNAs immobilized inside the fixed cells, excess unhybridized DOs are washed away and the remaining DOs, correctly hybridized to mRNA targets, are ligated with a DNA ligase. The *in situ* hybridized cells with ligated DOs still bound to target mRNAs are then transferred to a 96 well plate containing 48 unique first round barcodes to begin the combinatorial split pooling process that generates barcode diversity to uniquely label DOs within cells. The first-round barcodes, added by a second ligation to the previously hybridized and ligated DOs, have the advantage of easily marking sample identity before the cells are mixed in subsequent rounds of barcoding. The split pool process is repeated two more times to add the second and third round barcodes by ligation.

**Figure 1.**
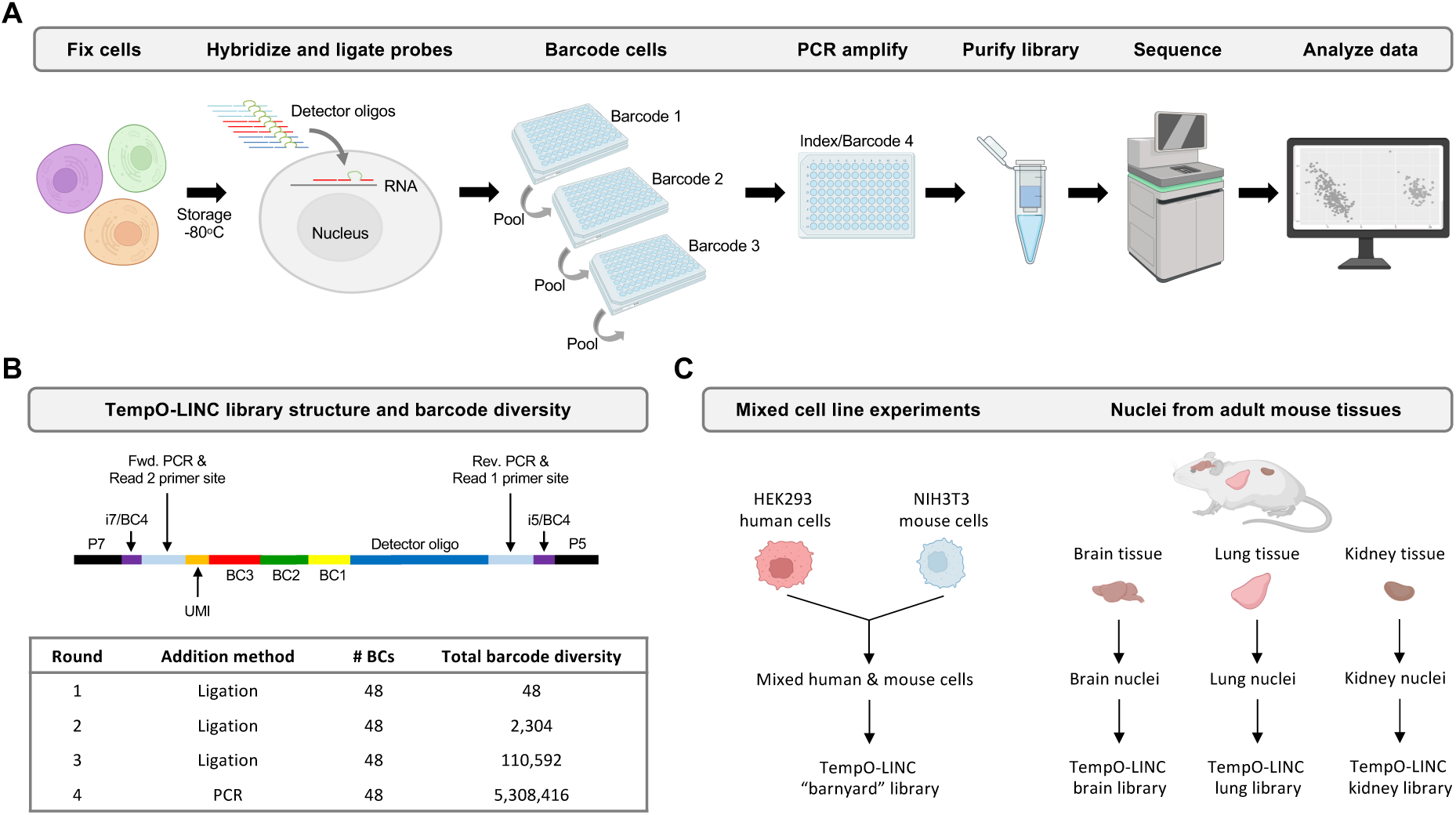
TempO-LINC workflow and study overview. (A) The TempO-LINC workflow begins with fixation of cells or nuclei and subsequent storage at -80°C. Fixed cells are thawed and hybridized with Detector Oligo (DO) probe pools that are then ligated. Split pool barcoding with three rounds of ligation is then performed. Following ligation of barcodes, cells are pooled again and redistributed into a final 96-well plate where sequencing adaptors and index sequences are added during PCR amplification. Amplified libraries are then purified and sequenced. Data analysis begins with demultiplexing to associate gene expression profiles with cell-identifying barcodes. (B) Schematic of TempO-LINC 310 base pair library structure including the three barcodes that are ligated as well as the i5 and i7 index positions that serve as a fourth barcode to identify cells. The table shows total barcode diversity created through each round of barcoding as well as the molecular mechanism for barcoding. (C) Overview of TempO-LINC validation experiments performed for this study. Experiments were a combination of gene expression profiling on mixtures of multiple human and mouse cell lines as well as nuclei isolated from adult mouse brain, lung and kidney tissue.

Although workflows for single-cell RNA sequencing can be laborious, our unique probe-based approach significantly simplifies sequencing library preparation. After the three rounds of split pool barcoding, intact cells are directly transferred to PCR reactions where the barcoded DOs are amplified, and the sequencing adaptors are added (Figure 1A). Following amplification, the DNA is spin-column purified and quantified to prepare the library for sequencing. Because each barcoded DO already has a forward and reverse priming site for the PCR and all barcoded DOs are the same length, there is no need for time consuming second strand cDNA synthesis or size fragmentation/selection steps necessary with traditional reverse transcription-based single-cell RNA-Seq approaches (Figure 1B).

The TempO-LINC approach is easily scalable based on the number of barcodes and rounds of barcoding performed. For the experiments reported in this study, we used three rounds of 48 barcodes that are ligated to the DOs as well as a fourth round of barcoding carried out during the amplification step with PCR primers using up to 48 unique dual indices (UDIs) (Figure 1B). The current format provides a maximum barcode diversity of 5,308,416 total barcode combinations.

To validate the performance of TempO-LINC, we identified several sample types to investigate, as illustrated in Figure 1C. The first set of experiments were performed on cell line samples consisting of a mixture of either human HEK293T and mouse NIH3T3 cells or four different human cell lines (MCF7, K562, HepG2 and Raji cells). Additionally, we chose to investigate complex mouse brain, lung and kidney tissues to demonstrate the ability of TempO-LINC to accurately profile and identify a variety of tissue-specific cell types. To demonstrate the performance of TempO-LINC on single nuclei, a challenging sample type that is frequently used for single-cell studies, we used nuclei isolated from each of the mouse tissues for subsequent barcoding with the TempO-LINC workflow.

### Validation of TempO-LINC on human and mouse cell lines

Formaldehyde fixed human HEK293T and mouse NIH3T3 cells were labeled with either Human Whole Transcriptome v2.1 (22,533 DDO/UDO probe pairs, 19,683 genes) or Mouse Whole Transcriptome v1.1 (30,146 probe pairs, 21,400 genes) DO pools that were modified to contain fully phosphorylated DDOs (Trejo et al., 2019; Yeakley et al., 2017). Following ligation of the DDOs to the UDOs that were hybridized to each targeted RNA, equal numbers of human and mouse cells were mixed and barcoded with the TempO-LINC workflow. Sequencing libraries were generated for three independent replicate experiments performed on different days and replicate libraries were sequenced and demultiplexed to evaluate TempO-LINC performance. On average, the libraries had 86% (± 0.8%) of all reads correctly barcoded with an expected combination of the 4 barcodes. Read count-ranked barcode plots showed a distribution indicative of successful single-cell barcoding with a clear inflection point separating cell-associated barcodes from noise (Figure 2A). Barcodes ranking above the inflection point were informatically labeled as cells, resulting in a total of 59,066 cells identified from the three replicate barcoding experiments, with an average of 9,397 human HEK293T cells and 10,291 mouse NIH3T3 cells per experiment.

**Figure 2.**
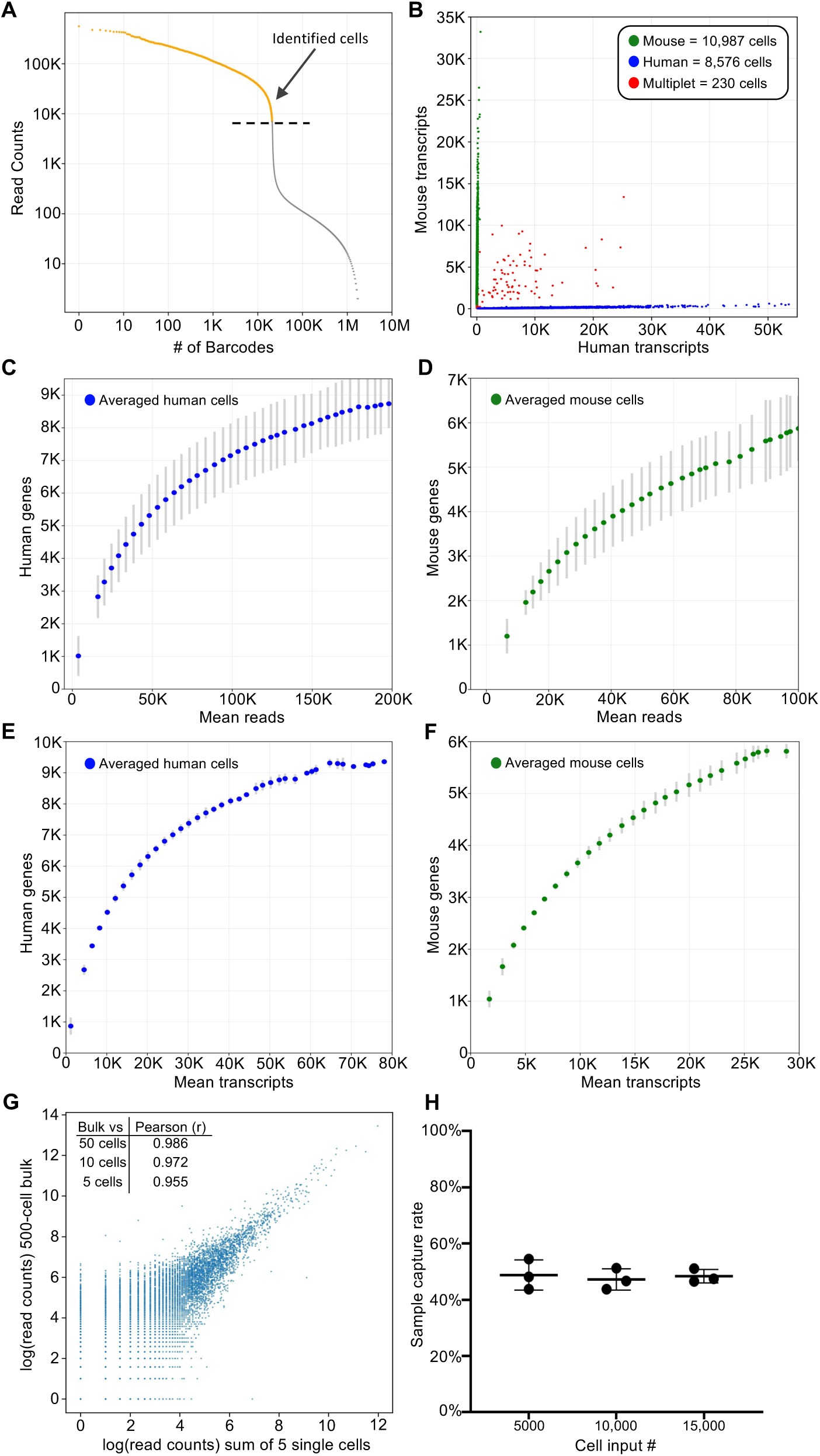
TempO-LINC enables high-throughput single-cell transcriptional profiling. Mixed human HEK293T and NIH3T3 cells were barcoded with TempO-LINC. Representative data from one of three replicates is presented in (A) and (B). (A) Read ranked barcode plot (knee plot) following demultiplexing of sequencing data. A clear inflection point is observed, above which, barcodes are identified as originating from a cell (shown in orange above the dashed line) (B) Plot showing cells with the number of transcripts mapping to either mouse or human genomes. Barcodes determined to be a multiplet consisting of both mouse and human cells are plotted in red. (C) Human or (D) mouse sequencing saturation curves showing the binned average number of genes detected at a given read depth for human HEK293T and mouse NIH3T3 cells from three replicate experiments with error bars representing the standard deviation. Data points represent the center value of 20,000 (human) or 25,000 (mouse) read bins. Average human (E) or mouse (F) genes detected per cell relative to increasing numbers of transcripts. Bins had a window size of 5,000 transcripts and are averaged across three experiments with standard deviation shown via error bars. (G) Correlation of read counts between a 500-cell bulk (not barcoded) sample and the sum/aggregate of read counts from 5 single cells expression profiled with TempO-LINC. The inset table also shows the Pearson (*r*) value from the correlation between the 500-cell bulk sample and the sum of either 50, 10 or 5 single HEK293T cells. (H) Sample capture rate plot showing the percentage of cells bioinformatically identified following the TempO-LINC assay. Three replicates were performed for varying initial sample cell input numbers of 5,000, 10,000 or 15,000 mouse NIH3T3 cells. Error bars indicate standard deviation.

Mixed species analysis was performed by plotting the number of mouse or human transcripts per cell (Figure 2B). From this analysis, an average multiplet rate of 0.54% (± 0.33%) was observed across the experiments at ∼20,000 cells analyzed per experiment. After adjusting the observed rates for non-detectable intraspecies multiplets and differing number of human and mouse cells, the actual TempO-LINC multiplet rate was calculated to be 1.09% (±0.66%) (Yamawaki et al., 2021).

Sequencing saturation curves for both mouse and human cells demonstrated sensitive and consistent gene detection rate performance across replicate TempO-LINC mixed species experiments. The averages of these replicate curves are plotted in Figure 2C-F. Average gene detection rates of cells within bins centered on 50,000 reads per cell, a common target for sequencing depth, were 5,311 genes/cell and 4,093 genes/cell for human and mouse cells, respectively. Transcript detection rates within these same bins were 14,605 transcripts/cell for human cells and 12,721 transcripts/cell for mouse cells. Due to the targeted nature of TempO-LINC, cells had low numbers of detected transcripts originating from mitochondrial (human and mouse <0.1%) or ribosomal genes

(human <4% and mouse <8%) and thus, the genes/cell are more likely to reflect actionable genes of interest. Additionally, analysis of the mRNA targeting position of the DOs demonstrates that they are broadly distributed across transcripts and lack the typical 3’ bias seen with traditional RNA sequencing approaches (Supplemental Figure 1).

To examine the potential impact of barcoding DOs on measurement of gene expression, we correlated the reads from a standard bulk assay (no barcoding but using the same set of DOs in the TempO-Seq assay) performed on 500 cells to aggregated reads from 50, 10 and 5 single cells identified with the TempO-LINC assay (Figure 2G). Even with only a 5-cell aggregate (pseudo-bulk), strong correlation was observed between bulk and single-cell assay gene expression (Pearson Correlation Coefficient (*r*) value of 0.955). To assess TempO-LINC assay reproducibility, we examined the correlation of human and mouse reads from replicate experiments and observed that the aggregate of reads from 50 human or mouse cells from each experiment were well correlated to each other (Pearson *r* values from 0.924 to 0.957, Supplemental Figure 2).

Efficient cell recovery or “capture” rates are critical for the identification of rare cell types within a population. We determined the cell recovery rates obtained from varying cell inputs into the TempO-LINC assay using 15,000, 10,000 or 5,000 fixed mouse NIH3T3 cells for the DO hybridization. Following DO ligation and barcoding with TempO-LINC, libraries were sequenced, and the number of cells recovered was determined by identifying barcodes above the inflection point in a ranked barcode plot. Consistent cell recovery rates of between 43.8% and 54.5% were observed across the replicates of all three input cell numbers with an overall average recovery rate of 48.2% (±3.6%) (Figure 2G).

To determine if TempO-LINC could transcriptionally profile and resolve mixtures of different cell types from the same species, we barcoded four human cell lines, MCF7, K562, HepG2 and Raji cells as well as a sample containing a mixture of all four cell lines together. Each independent cell line as well as the cell line mixture received unique first round barcodes within the same TempO-LINC experiment. Following sequencing and demultiplexing, we performed unsupervised clustering on all identified cells using Seurat (v5.0.3). Four distinct populations of cells were identified and displayed on a Uniform Manifold Approximation and Projection (UMAP) plot (Supplemental Figure 3). By identifying the cells within these clusters that were specific for one of the unique cell line sample barcodes, we observed that each UMAP cluster was composed of only one cell type, demonstrating the ability of TempO-LINC to accurately resolve multiple distinct human cell types from mixed populations based on whole transcriptome expression profiling.

### Single-nucleus Profiling of Adult Mouse Kidney

Having established a baseline for TempO-LINC performance on cell lines, we next sought to investigate how nuclei isolated from complex tissue performed in the assay. As a first test of this, nuclei from 45 mg of flash frozen adult mouse kidney tissue were isolated and then fixed with the TempO-LINC Fixation Kit. The nuclei were subsequently barcoded and sequenced with the standard TempO-LINC assay workflow using the mouse whole transcriptome DO pool. Following demultiplexing and analysis of the ranked barcode plot, 11,975 kidney cells were identified with an average read depth of 94,493 reads per cell (Figure 3A). Cells had mean transcript and gene counts of 5,649 transcripts per cell and 1,982 genes per cell, respectively (Figure 3A-C).

**Figure 3.**
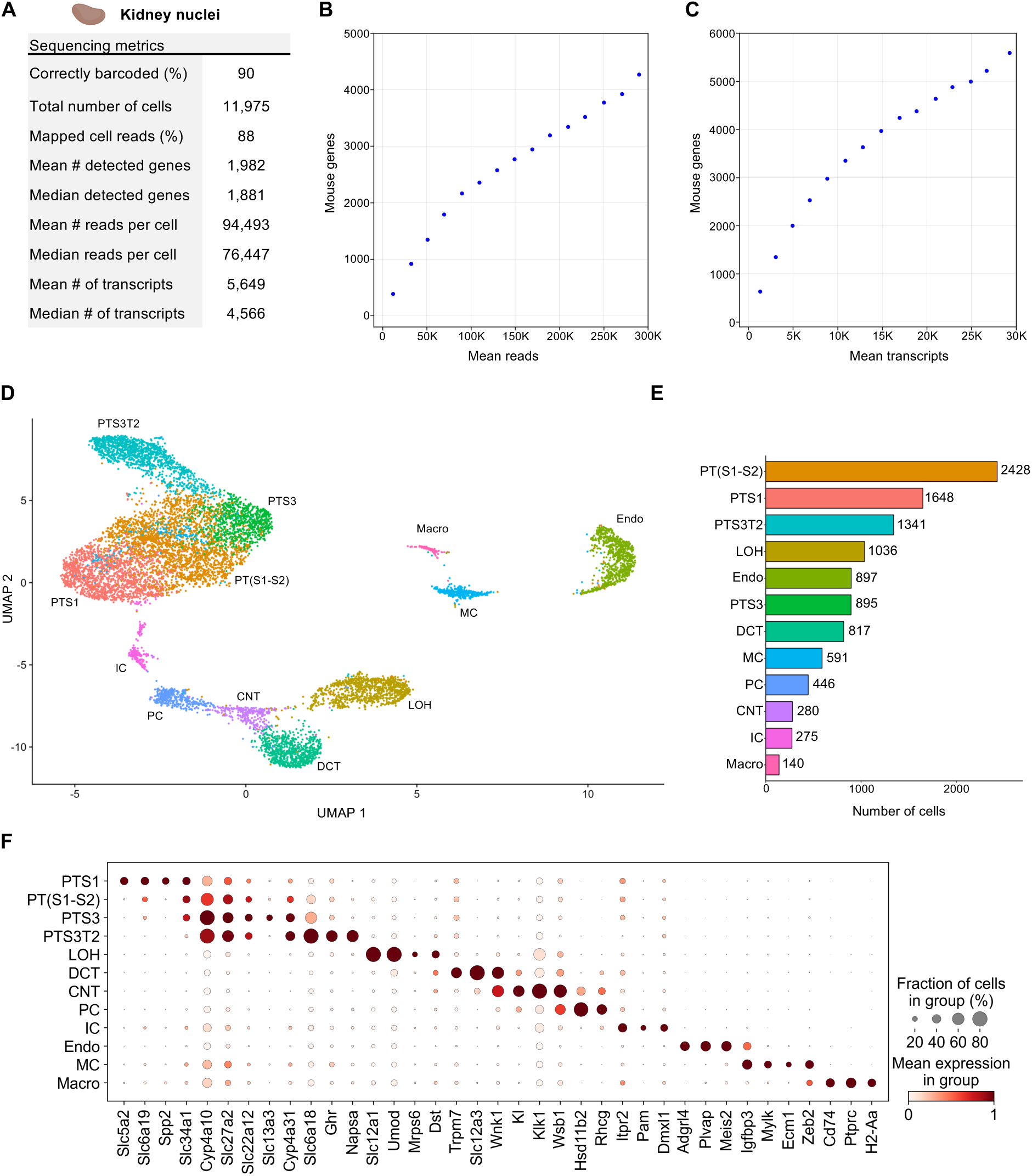
Single Nuclei Profiling of Adult Mouse Kidney. Nuclei from adult mouse kidney tissue were barcoded with TempO-LINC. (A) Table of key performance metrics following sequencing and demultiplexing of kidney nuclei libraries. (B,C) Sequencing saturation curves showing average number of genes detected per cell with increasing read or transcript depth. (D) 10,974 cells were plotted on a UMAP following unsupervised clustering. 12 distinct populations were identified and annotated using data from the MKA: proximal tubule segment 1 (PTS1), a mixture of proximal tubule segments 1 and 2 (PT(S1-S2), proximal tubule segment 3 (PTS3), proximal tubule segment 3 type 2 (PTS3T2), collecting duct intercalated cells (IC), principal cells (PC), connecting tubule (CNT), distal convoluted tubule (DCT), loop of Henle (LOH), mesangial cells (MC), endothelial cells (Endo) and macrophages (Macro). (E) Bar plot graph showing the number of cells within each population of clustered cells. (F) The expression level of representative kidney cell type biomarkers was examined for each of the 12 populations identified following unsupervised clustering.

Seurat was used to filter out low quality cells and perform unsupervised clustering on 10,794 mouse kidney cells identified with TempO-LINC. Visualization of cell clusters on a UMAP plot identified at least 12 distinct kidney cell populations (Figure 3D,E). We annotated these UMAP clusters based on their differential gene expression using the adult mouse kidney atlas (MKA) (Novella-Rausell et al., 2023). The largest class of cells consisted of proximal tubule (PT) cells which could be further classified into 4 distinct cell type clusters representing different segments of the tubule – segments 1, a mixture of segments 1 and 2, segment 3 and segment 3 type 2 (PTS1, PT(S1-S2), PTS3 and PTS3T2). Other cell clusters successfully annotated with the MKA included collecting duct intercalated cells (IC), principal cells (PC), connecting tubule (CNT), distal convoluted tubule (DCT), loop of Henle (LOH), mesangial cells (MC), endothelial cells (Endo) and macrophages (Macro). To further confirm our MKA-based cluster annotation and visualize known kidney specific biomarker expression within the assigned cell types, dot plots were generated showing the cluster-specific single-cell expression levels of 37 genes (Figure 3F).

One cell type that was not identified through unsupervised clustering was podocytes. Although these cells are often not identified with single-cell sequencing, it has been reported that sequencing of kidney nuclei preparations display better representation of podocytes (H. Wu et al., 2019). We therefore investigated whether any podocytes could be identified independently of clustering or if this cell type was completely absent from the data set. To identify podocytes from the 10,794 nuclei dataset, we performed overrepresentation analysis based on a defined gene set comprised of 7 well-known podocyte biomarkers - Nphs1, Nphs2, Podxl, Wt1, Magi2, Synpo and Thsd7a (Supplemental Figure 4) (Karp et al., 2021; M. C. Wu & Lin, 2009). This approach enabled the statistically significant identification of 7 podocyte cells within our single-nuclei dataset (false discovery rate (FDR) <0.0005). Identification of podocytes within this dataset indicates that, at least when looking for known transcriptional signatures, TempO-LINC has the sensitivity to identify rare cells comprising only ∼0.06% of the total population. We hypothesize that sequencing a larger number of nuclei would identify additional podocytes and enable them to be grouped together via unsupervised clustering.

### Single-nucleus Profiling of Adult Mouse Brain

Nuclei from 65 mg of flash frozen adult mouse brain tissue were isolated and then fixed with the TempO-LINC Fixation Kit. After fixation and subsequent barcoding with the TempO-LINC combinatorial workflow, brain libraries were sequenced and demultiplexed. Analysis of the ranked barcode plot identified 10,168 brain cells with an average read depth of 90,392 reads per cell (Figure 4A). Cells had mean transcript and gene counts of 9,017 transcripts per cell and 2,895 genes per cell, respectively (Figure 4A-C).

**Figure 4.**
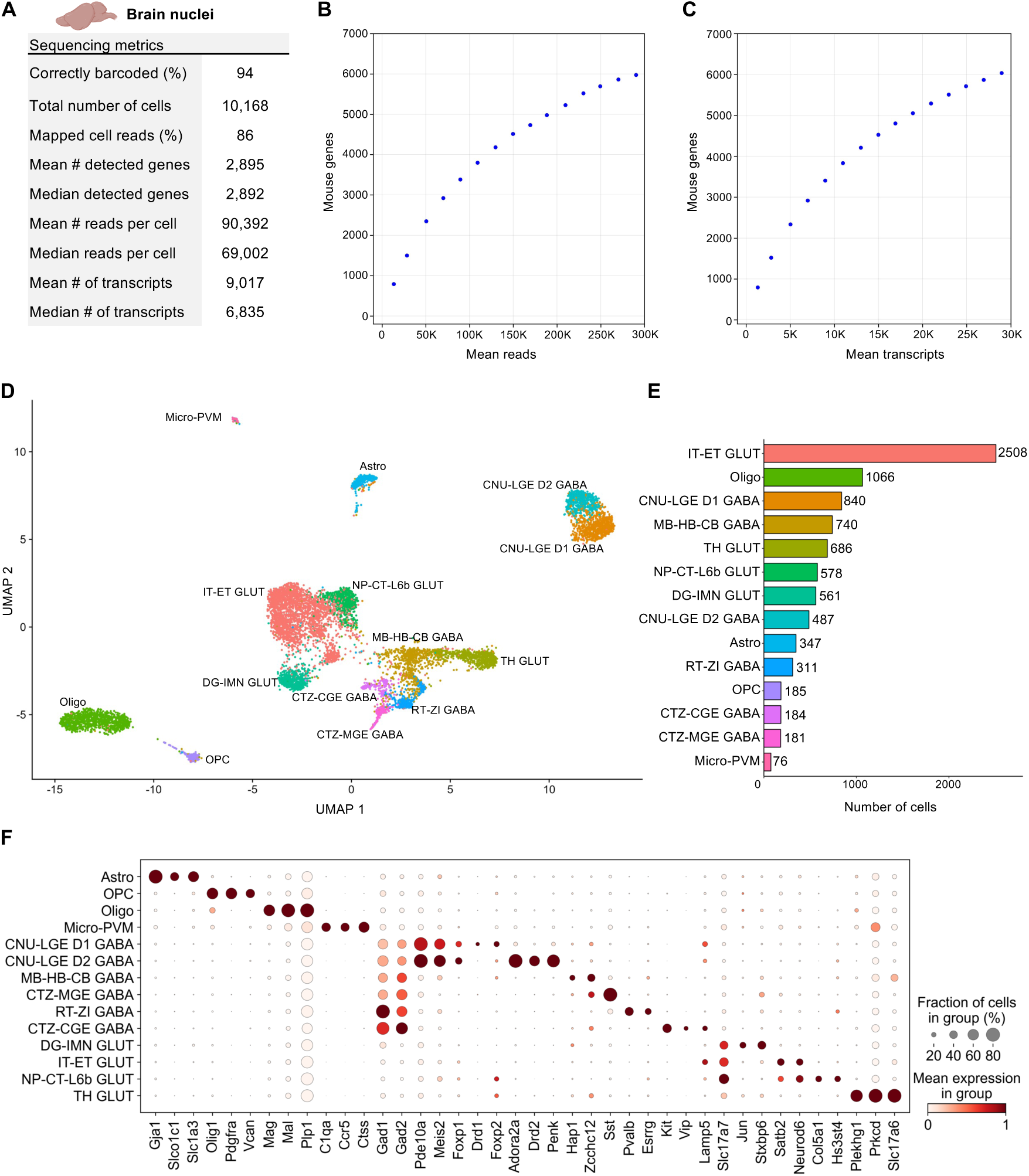
Single Nuclei Profiling of Adult Mouse Brain. Nuclei from adult mouse brain tissue were barcoded with TempO-LINC. (A) Table of key performance metrics following sequencing and demultiplexing of brain nuclei libraries. (B,C) Sequencing saturation curves showing average number of genes detected per cell with increasing read or transcript depth. (D) 8,750 cells were plotted on a UMAP following unsupervised clustering. 14 distinct populations were identified and annotated using data from the Allen Brain Institute (ABI): perivascular macrophages (micro-PVM), astrocytes (Astro), oligodendrocytes (Oligo), oligodendrocyte precursor cells (OPC), MB-HB-CB GABA neurons, RT-ZI GABA neurons, CTZ-CGE GABA neurons, CTZ-MGE GABA neurons, CNU-LGE D1 GABA neurons, CNU-LGE D2 GABA neurons, IT-ET GLUT neurons, NP-CT-L6b GLUT neurons, TH GLUT neurons and DG-IMN GLUT neurons. (E) Bar plot graph showing the number of cells within each population of clustered cells. (F) The expression level of representative braincell type biomarkers was examined for each of the 14 cell classes identified following unsupervised clustering. Gad1 and Gad2 are seen broadly expressed across GABA neurons as would be expected based on their molecular function. Similarly, Slc17a7 and Slc17a6 (human VGLUT1 and VGLUT2) are implicated in glutamate transport and mark multiple populations of glutamatergic neurons.

Following filtering in Seurat, the remaining 8,750 high-quality cells were clustered in an unsupervised manner based on differential gene expression and plotted as a UMAP (Figure 4D). At least 14 distinct populations were identified from the Seurat clustering. The Allen Brain Institute’s (ABI) Map My Cells automated annotation tool was used to perform initial identification and classification of the 14 cell clusters (Hao et al., 2021; Yao et al., 2023; Yao, Liu, et al., 2021). Four non-neuronal cell types – perivascular macrophages (micro-PVM), astrocytes (Astro), oligodendrocytes (Oligo), oligodendrocyte precursor cells (OPC) – and two major classes of neurons – GABAergic and glutamatergic – were identified. GABAergic neurons could be further subdivided into 5 classes – MB-HB-CB GABA, RT-ZI GABA, CTZ-CGE GABA, CTZ-MGE GABA and CNU-LGE GABA neurons. For glutamatergic neurons, 4 classes were identified – IT-ET GLUT, NP-CT-L6b GLUT, TH GLUT and DG-IMN GLUT. The number of cells populating each UMAP cluster are shown in Figure 5E. Following initial annotation with the Map My Cells tool, marker genes used by the ABI as well as single-cell brain studies were used to generate cluster specific dot plots and confirm the expected annotated cell type-specific expression of 40 biomarker genes across the brain populations (Figure 4F) (Cheng et al., 2022; Tasic et al., 2016; Yao et al., 2023; Yao, van Velthoven, et al., 2021).

**Figure 5.**
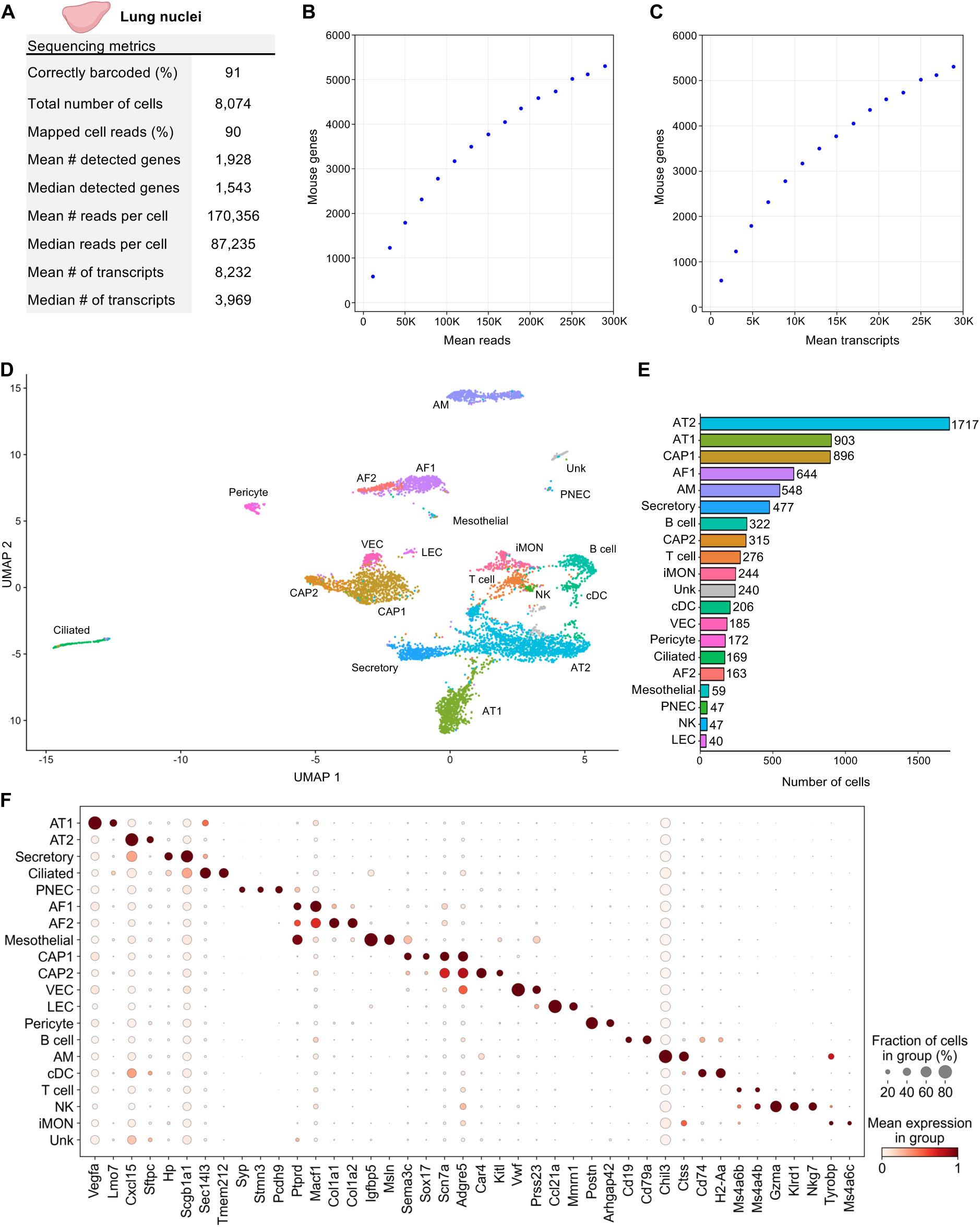
Single Nuclei Profiling of Adult Mouse Lung. Nuclei from 8-week-old adult mouse lung tissue were barcoded with TempO-LINC. (A) Table of key performance metrics following sequencing and demultiplexing of lung nuclei libraries. (B,C) Sequencing saturation curves showing average number of genes detected per cell with increasing read or transcript depth. (D) 7,670 cells were plotted on a UMAP following unsupervised clustering. A total of 20 distinct populations were identified and annotated using data from the mouse LungMAP Consortium, available through the Lung Gene Expression Analysis (LGEA) web portal: alveolar type 1 (AT1), alveolar type 2 (AT2), capillary 1 (CAP1), capillary 2 (CAP2), venous endothelial cell (VEC), alveolar fibroblast 1 (AF1), alveolar fibroblast 2 (AF2), Secretory, Ciliated, Pericyte, alveolar macrophage (AM), B cell, T cell, inflammatory monocytes (iMON), classical dendritic cell (cDC), mesothelial (Mesothelial), natural killer (NK), lymphatic endothelial cell (LEC), pulmonary neuroendocrine cell (PNEC) and an unknown population (Unk). (E) Bar plot graph showing the number of cells within each population of clustered cells. (F) The expression level of 42 representative lung cell type biomarkers was examined for each of the 20 populations identified following unsupervised clustering.

Although major classes of neurons are shown in Figure 5, increasing the clustering resolution within Seurat and/or feature plotting specific biomarkers could delineate additional subclasses of neurons. One example of this is shown for the CNU-LGE GABA class of neurons that could be further divided into two subclasses on the UMAP (Figure 4D). Both D1 and D2 dopamine receptor expressing spiny neuron subtypes were identified within the CNU-LGE GABA population, CNU-LGE D1 GABA and CNU-LGE D2 GABA, respectively (Anderson et al., 2023; He et al., 2021; Zywitza et al., 2018). Similarly, we demonstrated that 5 additional neuronal subclasses could be found within the IT-ET GLUT population following increasing the resolution on this cluster in Seurat (Supplemental Figure 5). Using the ABI automated annotation tool combined with neuronal subtype specific gene expression analysis, these subclusters were identified as predominantly L2/L3 IT, L4/L5 IT, L6 IT, L5 ET and Ca1-ProS cortical neurons (Cheng et al., 2022; Tasic et al., 2016; Yao et al., 2023; Yao, van Velthoven, et al., 2021). The identification of these additional cell types within the IT-ET GLUT population brought the total number of distinct brain cell types identified to 18. A similar strategy of increasing the resolution on other clusters would likely increase this number further but was beyond the scope of our current study. Nevertheless, our results indicate that TempO-LINC can generate rich datasets displaying a diverse range of known cell types from tissues.

### Single-nucleus Profiling of Adult Mouse Lung

Nuclei from 50 mg of flash frozen adult mouse lung tissue were isolated and then fixed with the TempO-LINC Fixation Kit. The fixed nuclei were then barcoded and sequenced with the TempO-LINC workflow. Following demultiplexing and analysis of the ranked barcode plot, 8,074 lung cells were identified with an average read depth of 170,356 reads per cell (Figure 5A-C). Cells had mean transcript and gene counts of 8,232 transcripts per cell and 1,928 genes per cell, respectively.

As with the datasets from kidney and brain tissue, lung cells were further filtered in Seurat and 7,670 high-quality cells were used for unsupervised clustering. A total of 20 clusters were visualized on the UMAP and subsequently annotated using mouse LungMAP Consortium data, available through the Lung Gene Expression Analysis (LGEA) web portal, as a guide (Figure 5D) (M. Guo et al., 2023; Sun et al., 2022). Major classes of cells represented across the 20 identified populations include endothelial (CAP1, CAP2, VEC LEC), mesothelial, epithelial (AT1, AT2, Secretory, Ciliated, PNEC), fibroblast (AF1, AF2) pericyte, and immune (AM, B cell, T cell, cDC, iMON, NK) cell types. There was also one unknown (Unk) cell population that we were unable to confidently annotate based on the available lung reference data markers. Notably, we were able to utilize unsupervised clustering to identify four cell types that each represented between 0.8% and 0.5% of the total population, including known rare cell types such as PNECs (Figure 5E) (Kuo et al., 2022). Figure 5F shows the cluster specific expression of 42 lung cell type marker genes (M. Guo et al., 2023; Hurskainen et al., 2021).

## Discussion

The presented data demonstrate that TempO-LINC is a robust, instrument-free platform for high-throughput single-cell gene expression profiling. The approach is fundamentally different than almost all other single-cell expression profiling methods in that it does not rely on reverse transcription of mRNA. Of the commercially available solutions for single-cell expression profiling, only the Chromium Fixed RNA Profiling (FRP) kit from 10X Genomics has a similar approach to TempO-LINC. The Chromium FRP kit relies on an oligonucleotide ligation assay (OLA) and was recently shown to display greater sensitivity than reverse transcription-based methods, albeit at a cost far greater than the combinatorial split pool approaches evaluated (De Simone et al., 2024). With TempO-LINC, we introduce a novel method that combines oligonucleotide ligation-based chemistry with highly scalable combinatorial barcoding that does not rely on expensive instrumentation or emulsion assays.

High single-cell gene detection rates are critical for obtaining maximal biological insight into cell states and behaviors. We observed both high sensitivity barcoding and gene detection rates with TempO-LINC. Sequencing libraries across our study had between 85% and 94% correctly barcoded reads and, although it is not possible to directly compare gene detection rates across platforms without costly benchmarking experiments, our observed rates of 5,311 human genes/cell and 4,156 mouse genes/cell at a depth of ∼50,000 reads/cell compares favorably to reports of the best performing high-throughput single-cell expression assays from previous studies (Ding et al., 2020; Yamawaki et al., 2021). Additionally, the targeted nature of the TempO-LINC approach resulted in a low percentage of reads occupied with mitochondrial and ribosomal transcripts. This effectively increases the sensitivity of TempO-LINC and likelihood that the approach will identify potentially more informative genes expressed at lower levels. Similarly, attenuation is another feature of TempO-LINC that further increases gene detection sensitivity by using non-functional competitor DOs to reduce the sequencing space/read depth occupied by specific highly expressed genes, thereby increasing the dynamic range and gene detection rate of the assay. In total, 3.86% (869 probe pairs) of Human Whole Transcriptome v2.1 and 1.76% (531 probe pairs) of Mouse Whole Transcriptome v1.1 DO pools were attenuated in this study. Attenuation factors range between 2 and 49-fold depending on the specific probe pair. Importantly, attenuation reduces, but does not eliminate the read counts from the attenuated genes and has been shown to have no impact on the ability to perform differential expression analysis on these genes (Yeakley et al., 2017). The net result of both attenuation and low ribosomal and mitochondrial gene targeting is that for a given number of transcripts detected within a cell, TempO-LINC displays higher gene detection rates than other methods (e.g., >6,000 genes detected at 20,000 transcripts, Figure 2E).

Even though we restricted this study to transcriptome-wide analysis of human and mouse cells, smaller probe sets can be chosen to target select genes and pathways. For example, the TempO-Seq S1500+ Surrogate Assay DO pool targets only ∼2,200 RefSeq genes and can be used to cost-effectively monitor transcriptional perturbation of major molecular pathways involved in toxicological responses (Bushel et al., 2018; Mav et al., 2018). This panel or even smaller custom panels are compatible with the TempO-LINC approach and would reduce the number of reads required to reach sequencing saturation. Moreover, as we demonstrated, oligonucleotide probes can be targeted to any region within mRNAs; consequently, the assay does not have the 3’ bias inherent to transcriptomic analysis with common oligo-dT priming strategies. Probes can also be targeted to specifically detect gene fusions, variants, and/or alternative splicing events at almost any splicing junction, enabling panels for monitoring of splicing at single-cell resolution (Oh et al., 2021; Yeakley et al., 2002).

Cell capture efficiency is a key metric for single-cell expression profiling with higher capture rates enabling optimal performance on samples having few cells or important rare cell populations. These rates can vary substantially across methods with ddSEQ and Drop-seq methods capturing <2% of cells and the Chromium (10X Genomics) platform having rates from ∼30% to ∼80% (Yamawaki et al., 2021). The TempO-LINC capture rate of ∼50% is therefore one of the highest reported rates and makes the method well-suited for studying samples where expression profiling the maximum number of cells within the sample is a priority. Another critical metric to consider for accurate single-cell expression profiling is the multiplet or doublet rate. Multiplet rates have been reported around 2-3% for split pool barcoding approaches and ∼2-8% with droplet-based Chromium (Ding et al., 2020; Hornung et al., 2023; Lareau et al., 2020; Yamawaki et al., 2021; Zheng et al., 2017). With TempO-LINC, the multiplet rate was <1.1% for ∼20,000 cells detected, placing this method amongst the lowest reported multiplet rates of any platform.

Many clinically important studies rely on formalin-fixed paraffin-embedded (FFPE) samples. Despite the potential insight and utility that could be gained with single-cell RNA sequencing of these samples, the RNA degradation and chemical modification within FFPE derived cells severely impacts the performance of single-cell expression profiling methods that require cDNA synthesis. It has recently been reported that targeted probe-based approaches enable improved single-cell data quality from FFPE samples relative to conventional scRNA-Seq with reverse transcription (Y. Guo et al., 2023; Vallejo et al., 2022). Given TempO-Seq also uses short, targeted probes and can produce high-quality gene expression data from bulk FFPE samples, we anticipate TempO-LINC will similarly enable single-cell resolution from FFPE isolated cells and/or nuclei (Cannizzo et al., 2022; Trejo et al., 2019).

From this study, we conclude that TempO-LINC is a novel, instrument-free, single-cell genomics platform capable of resolving complex cellular heterogeneity from a variety of sample types including nuclei. We continue to expand the validated applications and samples demonstrated with the assay. Moreover, the foundational targeted DO approach has also been implemented for detection of genetic variants, epigenetic markers and antibodies; consequently, high-throughput single-cell multi-omic assays are additional focus areas for TempO-LINC development.

## Materials and Methods

### Cell culture and sample preparation

Cryopreserved vials of cells were obtained from American Type Culture Collection (ATCC). HEK293T (ATCC, cat# CRL-1573) and NIH3T3 (ATCC, cat# CRL-1688) cells were thawed at room temperature for 5-10 minutes and cultured for a minimum of three days before proceeding to the TempO-LINC fixation protocol. NIH3T3 cells were cultured in Dulbecco’s Modification of Eagle’s Medium (DMEM; Corning, cat# 10-013-CM) with heat-inactivated 10% Fetal Bovine Serum (FBS; Omega Scientific, FB-02), and HEK293T cells were cultured in Roswell Park Memorial Institute (RPMI; Gibco, 11875-093) 1640 with heat-inactivated 10% FBS media. Prior to cell fixation, cells were cultured at 37°C and 5% CO2. Once cells reached ∼80% confluency, cells were collected using 0.05% Trypsin-EDTA (Gibco, cat# 25300-054), and collected in 15 mL conical tubes. Trypsin-EDTA was neutralized with DMEM + 10% FBS or RPMI 1640 + 10% FBS for NIH3T3 or HEK293T cells, respectively. Cells were pelleted via swinging-bucket centrifugation for 5 minutes at 400 rcf and 4°C. The supernatant was aspirated, and the pellet was resuspended in 5-10 mL of pre-warmed media to produce a single-cell suspension. Cells were quantified using a TC20 Automated Cell Counter (Bio-Rad, cat# 1450102) and briefly stored at 4°C.

For the mixed human cell line experiment, cell lines used were MCF7 (ATCC, cat# HTB-22), HepG2 (ATCC, cat# HB-8065), Raji (ATCC, cat# CCL-86) and K562-r (ATCC, cat# CRL-3344). Cells were cultured according to ATCC recommendations and fixed with the TempO-LINC fixation protocol. Each cell line was placed into 4 wells of the TempO-LINC Barcode 1 plate while the mixed cell line population consisted of roughly equal numbers of each fixed cell line that were combined and barcoded with 32 unique round 1 barcodes. Barcoding and sequencing were carried out as described below.

### Nuclei Preparation

Fresh tissue was isolated from an 8-week-old ICR (CD-1^®^) mouse following IACUC-approved protocols. The harvested tissues included one cerebral cortex hemisphere (65 mg), one half of a kidney (45 mg), and one lung lobe (50 mg), which were immediately snap-frozen on dry ice. The frozen tissues, along with 75 µL of RNase Inhibitor V2 (S2 Genomics, 100-288-916), were then placed in the Dissociation Chamber of a pre-cooled standard Nuclei Isolation Cartridge (S2 Genomics, 100-063-287) and inserted into the pre-cooled Singulator 100^TM^ (S2 Genomics, 100-060-817). Nuclei were isolated from the tissues using the Low Volume Nuclei Isolation protocol.

Following isolation, the nuclei suspension was recovered from the Singulator cartridge and centrifuged at 500 rcf for 5 minutes at 4°C. The supernatant was removed, and the kidney and lung nuclei pellets were resuspended in 1 mL of Nuclei Storage Reagent (S2 Genomics, 100-063-405) supplemented with RNase inhibitor (1 U/µL). The brain nuclei were further processed using Debris Removal Stock Reagent (S2 Genomics 100-253-628) to eliminate myelin debris, according to the manufacturer’s instructions. Finally, the nuclei were counted, and their viability was determined using AO/PI staining with the Nexcelom K2. The nuclei samples were subsequently fixed via the TempO-LINC Fixation protocol as described below, and stored at -80°C.

### TempO-LINC fixation protocol

Cells were fixed using the TempO-LINC Fixation Kit (BioSpyder Technologies Inc., cat# 202392). We started with 100,000 – 400,000 high percent viability (> 90%) cells, with minimal cell clumps and doublets present (< 20%). Cells or nuclei were pelleted via centrifugation for 5 minutes at 400 rcf and 4°C, the supernatant was aspirated, and the pellets were resuspended in TempO-LINC Fixation Wash Buffer. During the spin, TempO-LINC Fixation Buffer was prepared using 16% methanol-free formaldehyde (Thermo ScientificTM PierceTM, cat# PI28906). Following the wash step, samples were pelleted and resuspended in TempO-LINC Fixation Buffer. Samples were then fixed for 1 hour at 25°C without agitation. Following fixation, samples were quenched with TempO-LINC Fixation Quenching Buffer. Samples were quantified and dispersed into 1.5 mL screw-cap tubes containing TempO-LINC Freezing Buffer. Tubes were placed in a freezing container (-1°C/minute cooling rate) and stored at -80°C until needed.

### Expression probe labeling and barcoding

TempO-LINC mixed species barcoding experiments were performed with a 50:50 split of human HEK293T and mouse NIH3T3 cells. Cryopreserved vials of fixed HEK293T and NIH3T3 cells were thawed at room temperature and centrifuged using a swinging-bucket centrifuge for 5 minutes at 500 rcf and 25°C. Each cell pellet was separately resuspended in 1 mL of TempO-LINC 1X Pre-Hybridization Buffer and quantified using the TC20 Automated Cell Counter. HEK293T cells and NIH3T3 cells were transferred to separate 1.7 mL microcentrifuge tubes and centrifuged for 5 min at 500 rcf. Both tubes of cells were resuspended in 55 µL TempO-LINC 1X Hybridization buffer. HEK293T cells were incubated with 45 µL of a Human Whole Transcriptome v2.1 (22,533 probe pairs, BioSpyder, cat #201828) detector oligo (DO) pool, and NIH3T3 cells were incubated with 45 µL of a Mouse Whole Transcriptome v1.1 (30,146 probe pairs, BioSpyder, cat# 201186) DO pool overnight for 16 hours at 45°C. Following hybridization, cells were centrifuged for 5 min at 500 rcf and were washed twice with 750 µL of 1X Ligation buffer. HEK293T and NIH3T3 cells were separately incubated with DNA Ligase mix for 1 hour at 37°C. Following DO ligation, cells were washed twice with 1X Ligation buffer and quantified.

Next, 50,000 HEK293T cells and 50,000 NIH3T3 cells were mixed in a single microcentrifuge tube for TempO-LINC barcoding. TempO-LINC Barcoding Ligation mix was prepared by supplementing 1X Barcoding Ligation buffer with TempO-LINC ligase and RNase Inhibitor. The combined HEK293T and NIH3T3 cell pellet was resuspended in 1225 µL of Barcoding Ligation mix and transferred to a polyvinyl chloride (PVC) reservoir (VWR cat# 89094-688). Using a multichannel pipette, cells were then distributed across 48 wells (2,083 cells/well) of the TempO-LINC Barcode 1 plate. The plate was sealed with an adhesive film and incubated for 30 minutes at 45°C using BioRad’s C1000 Touch Thermal Cycle/CFX96 Real-Time System (Bio-Rad, cat# 1845096). Following incubation, the plate was removed from the thermocycler, and 10 µL (per well) of TempO-LINC Barcode Blocker 1, were added across the Barcode 1 plate. The plate was sealed with an adhesive film and incubated for 10 minutes at 37°C. After incubation, all cells in the Barcode 1 plate were pooled, transferred to microcentrifuge tubes, and centrifuged for 5 minutes at 500 rcf at 25°C. The supernatant was aspirated and the cells resuspended in freshly prepared TempO-LINC Barcoding Ligation mix. Cells were then redistributed across 48 TempO-LINC Barcode 2 plate wells. The plate was incubated for 30 minutes at 45°C. Following barcode ligation, 10 µL of TempO-LINC Barcode Blocker 2, was added to each well of the Barcode 2 plate and incubated for 5 minutes at room temperature. After blocking, all cells in the Barcode 2 plate were pooled, transferred to microcentrifuge tubes, and centrifuged for 5 minutes at 500 rcf and 25°C. The supernatant was aspirated, and cells were resuspended in freshly prepared TempO-LINC Barcoding Ligation mix. Cells were redistributed across a TempO-LINC Barcode 3 plate. The plate was incubated for 30 minutes at 45°C. Following barcode ligation, 10 µL of TempO-LINC Barcode Blocker 3, was added to each well of the Barcode 3 plate and incubated for 5 minutes at room temperature. After barcode blocking, all cells were pooled, transferred to microcentrifuge tubes, and centrifuged for 5 minutes at 500 rcf and 25°C. The supernatant was aspirated, and cells were washed twice with 1X Barcode Wash buffer. The cell pellet was resuspended in 300 µL TempO-LINC PCR Amplification buffer.

### Barcoding of nuclei

Fixed, cryopreserved mouse nuclei were thawed at room temperature and centrifuged using a swinging-bucket centrifuge for 5 minutes at 500 rcf and 25°C. Nuclei were resuspended in 1mL of 1X TempO-LINC 1X Pre-Hybridization Buffer and quantified. 150,000 nuclei were transferred to 1.7 mL microcentrifuge tubes and centrifuged for 5 min at 500 rcf. Nuclei were resuspended in 55 µL TempO-LINC 1X Hybridization buffer and 45 µL of a Mouse Whole Transcriptome v1.1 (30,162 probe pairs, BioSpyder, cat# 201186) DO pool and placed in an incubator overnight for 16 hours at 45°C. Next, 100,000 nuclei were transferred and prepped for TempO-LINC Barcoding. Nuclei were equally distributed across the 48 wells of the TempO-LINC Barcode 1 plate at 2,083 nuclei/well nuclei and were barcoded identically to the cells as described previously. Following barcoding, nuclei were resuspended in 200 µL TempO-LINC PCR Amplification buffer.

### Barcoded library amplification and purification

Following the three rounds of ligation-based barcoding, cells or nuclei were then distributed for direct PCR amplification between 600-1200 cells/well or 1000 nuclei/well. The PCR was run using a C1000 Touch Thermal Cycle/ CFX96 Real-Time System. PCR primers contained both Illumina sequencing adapters as well as dual indices that also served as the fourth TempO-LINC barcode. PCR thermocycling conditions were as follows: incubation for 10 minutes at 37°C, initial denaturation for 3 minutes at 95°C, then the PCR cycled 28-30 times from 95°C for 15 seconds, to 65°C for 30 seconds, then 68°C for 30 seconds. Finally, the thermocycler briefly held temperature to 68°C for 2 minutes before permanently holding at 25°C.

TempO-LINC libraries were purified using the NucleoSpin Gel and PCR Clean-up Kit (Macherey-Nagel, cat# 740609.250), according to the manufacturer’s protocol. Double-stranded DNA-based TempO-LINC libraries were quantified using the DeNovix DS-11 Series spectrophotometer (DeNovix Inc., cat # DS-11) and the Qubit 4 Fluorometer (ThermoFisher Scientific, cat# Q33226) with the Qubit 1X dsDNA HS Assay Kit (ThemoFisher Scientific, cat# Q33231). All library quantification was performed according to the manufacturer’s protocol. TempO-LINC libraries were also assessed with the Agilent TapeStation 4150 (Agilent, cat# G2992A) and Agilent D1000 ScreenTape (Agilent, cat# 5067-5582), according to the manufacturer’s protocol. TempO-LINC library signal intensity was visualized using TapeStation Analysis Software v3.2 (Agilent).

### Sequencing and Demultiplexing

Libraries were sequenced on a NovaSeq 6000. Illumina S1 flow cells with v1.5 (200 cycle) reagent kits were used to sequence both the mixed species and nuclei experiments. For the mixed human cell line experiment, the library was sequenced on an Illumina MiniSeq using a Mid Output Kit (300 cycles).

To generate TempO-LINC single-cell expression data, barcoded reads are demultiplexed using an in-house developed software. Briefly, index associated reads are scanned to find the sequences of the barcodes, while allowing an offset of 3 nucleotides from their expected positions and 1 mismatch from their expected sequence. Reads featuring the same barcode combination are pooled together into barcode associated clusters. A cell calling read threshold is established based on inflection point analysis of a ranked barcode distribution plot (knee plot). Cell associated reads are then aligned to the reference human or mouse whole transcriptome TempO-Seq probe sequences using bwa v. 0.7.17 (mem algorithm, parameters: -v 1 -c 2 -L 100). Unique molecular identifiers (UMIs) are used to collapse the reads originating from the same transcript and remove any PCR duplicates. For each single-cell alignment, the probes featuring a UMI count >0 are considered indicative of an expressed/detected gene. Cell count tables are generated using the software featureCount v 2.0.1.

### Sequencing Saturation Analysis

To generate datapoints for the sequencing saturation curves for the mixed human and mouse species experiments, data from cells was binned and averaged. Binning was performed separately for each of the three experiments using a window size of 20,000 reads and a window step of 5,000 reads for human data, and a window size of 25,000 reads and a window step of 3,000 reads for mouse data. For each bin, cells with read counts within the bin range were collected, and the average number of detected genes and the average number of reads were calculated from these cells within each bin for each experiment. Next, the average number of detected genes across the three experiments was calculated for each bin, and the standard deviation of the detected genes was calculated. A scatter plot with error bars representing the standard deviation was then created to visualize the data. The mixed species transcript saturation curves were calculated in a similar manner; however, a window size of 5,000 transcripts and a window step of either 2,000 or 1,000 transcripts were used for human and mouse cell data, respectively.

Sequencing saturation curves for the nuclei datasets presented in Figures 3, 4 and 5 were also calculated by binning and averaging data from individual nuclei in each experiment. For the read-based saturation curves, a bin or window size of 20,000 reads and a window step of 20,000 were used. For transcript-based saturation curves, a window of 2,000 transcripts and a step size of 2,000 transcripts were used.

### Single-cell Clustering and Annotation

Single-cell mouse tissue datasets were processed with the R package Seurat (v5.0.3)(Hao et al., 2024). Briefly, genes with expression in less than 3 cells were excluded from the dataset matrix and low-quality cells were removed based on gene and read counts. The remaining high-quality cells were utilized for downstream analyses. Data were normalized using the relative-counts method with a scaling factor of 1 × 10^6^ within Seurat. The top 2000 highly variable genes were selected using the variance-stabilizing transformation approach and utilized in principal component analysis. Cells were then clustered using the K-nearest neighbor (KNN) approach followed by Louvain algorithm optimization and visualization using Uniform Manifold Approximation and Projection (UMAP). The number of cells per cluster were extracted using scCustomize (v2.1.2)(Marsh et al., 2021) and used to generate bar plots for each tissue dataset. Differential gene expression analysis was performed between clusters using a Wilcoxon rank-sum test and p-values were adjusted using the Bonferroni correction method for multiple-comparison. Cell clusters were annotated using a combination of differentially expressed markers genes and the R Azimuth annotation package (v0.5.0) (Hao et al., 2021), the LunGENS database(Ardini-Poleske et al., 2017; Du et al., 2015), and the Allen Brain Institute MapMyCells (Yao et al., 2023) annotation tools for kidney, brain, and lung tissues respectively. Dot plots denoting cluster marker genes were generated using scanpy (v1.10.1)(Wolf et al., 2018).

Subclustering of the mouse brain IT-ET GLUT population was performed by clustering solely the IT-ET GLUT using the subcluster function in Seurat. Similarly to whole brain cluster annotation, IT-ET GLUT subclusters were annotated using a combination of MapMyCells and marker genes identified via differential gene expression analysis.

Identification of podocytes within the mouse kidney was performed using overrepresentation analysis (ORA) (Beißbarth & Speed, 2004; Tavazoie et al., 1999) utilizing a seven podocyte marker gene set versus all genes present in the dataset. We considered any gene having greater than 150 normalized read counts as being positive for expression of one of these markers. ORA was conducted following a hypergeometric distribution (phyper R stats package v4.3.2)(R Core Team, 2023) and FDR value was calculated using the Benjamini & Hochberg adjustment method.

## Acknowledgements

The authors would like to kindly thank Nabiha Khan, Nate Pereira, Stevan Jovanovich and John Bashkin from S2 Genomics. S2 Genomics provided the Singulator 100 instrument and mouse tissue samples that were used in this study. Additionally, we thank Milos Babic and Nicole Martin for their assistance with early-stage development of the TempO-LINC workflow. This work was supported by NIH grants R43GM140771 and R44GM140771. Panels for some of the figures were created with BioRender.com.

## Author Contributions

D.J.E. managed the study, designed the TempO-LINC assay, planned the experiments to be performed, prepared nuclei for barcoding, analyzed the data and wrote the manuscript. K.S.W. prepared all reagents, performed the experiments, analyzed the data and edited the manuscript. N.D.J. carried out the data analysis (including bioinformatics and Seurat analysis of all the mouse nuclei samples) and edited the manuscript. S.C., H.H. and G.M. developed the barcode demultiplexing pipeline and analyzed the data. K.G.W. performed the mixed human cell line experiment. J.Y. assisted with primer design, sequence analysis and edited the manuscript. B.S. and J.M. conceived of the combinatorial approach for TempO-LINC and edited the manuscript.

B.S. is a PI on grants funding part of this work. All authors have read and approved this manuscript.

## Data Availability

Data from the reported experiments will be made available for viewing and download at https://www.biospyder.com/single-cell.

## Supplemental Figures

**Supplemental Figure 1.**
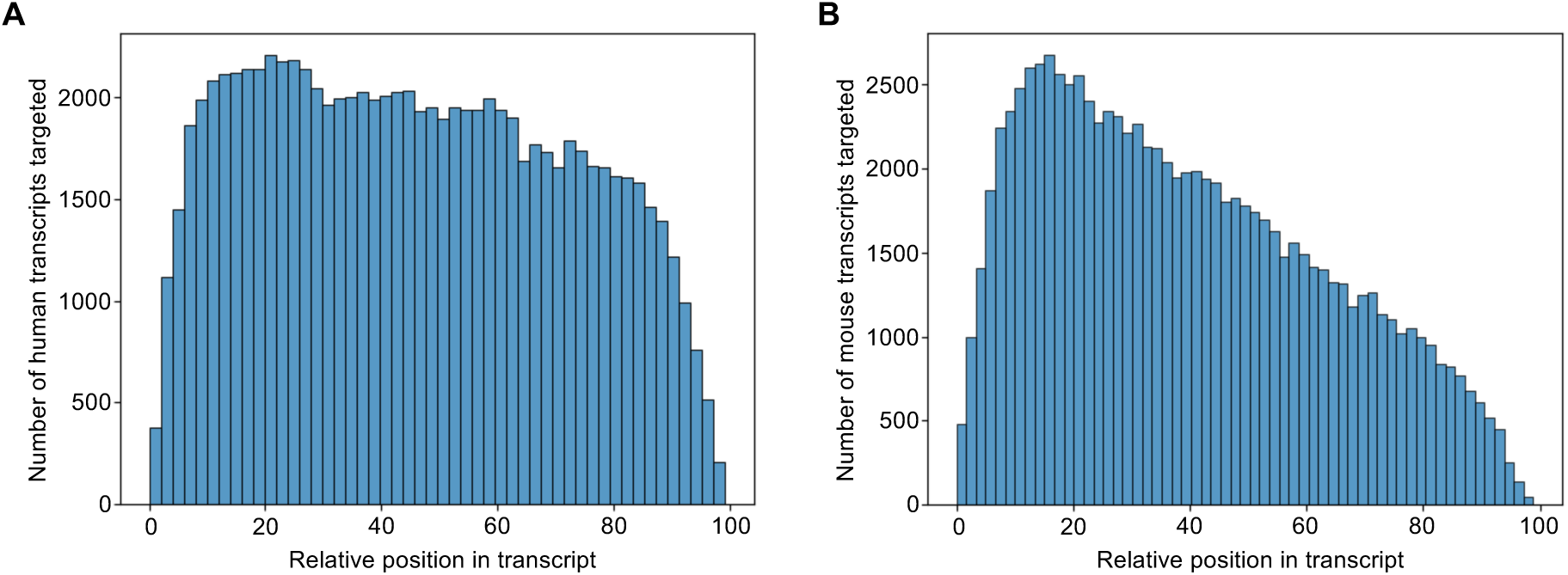
Bar plots showing the relative targeting position of DO probes within their target mRNAs with “0” representing the 5’ end and “100” the 3’ end of the message. (A) Analysis of the Human Whole Transcriptome v2.1 (22,533 probe pairs, 19,683 genes) DO pool. (B) Distribution of the Mouse Whole Transcriptome v1.1 (30,146 probe pairs, 21,400 genes) DO pool.

**Supplemental Figure 2.**
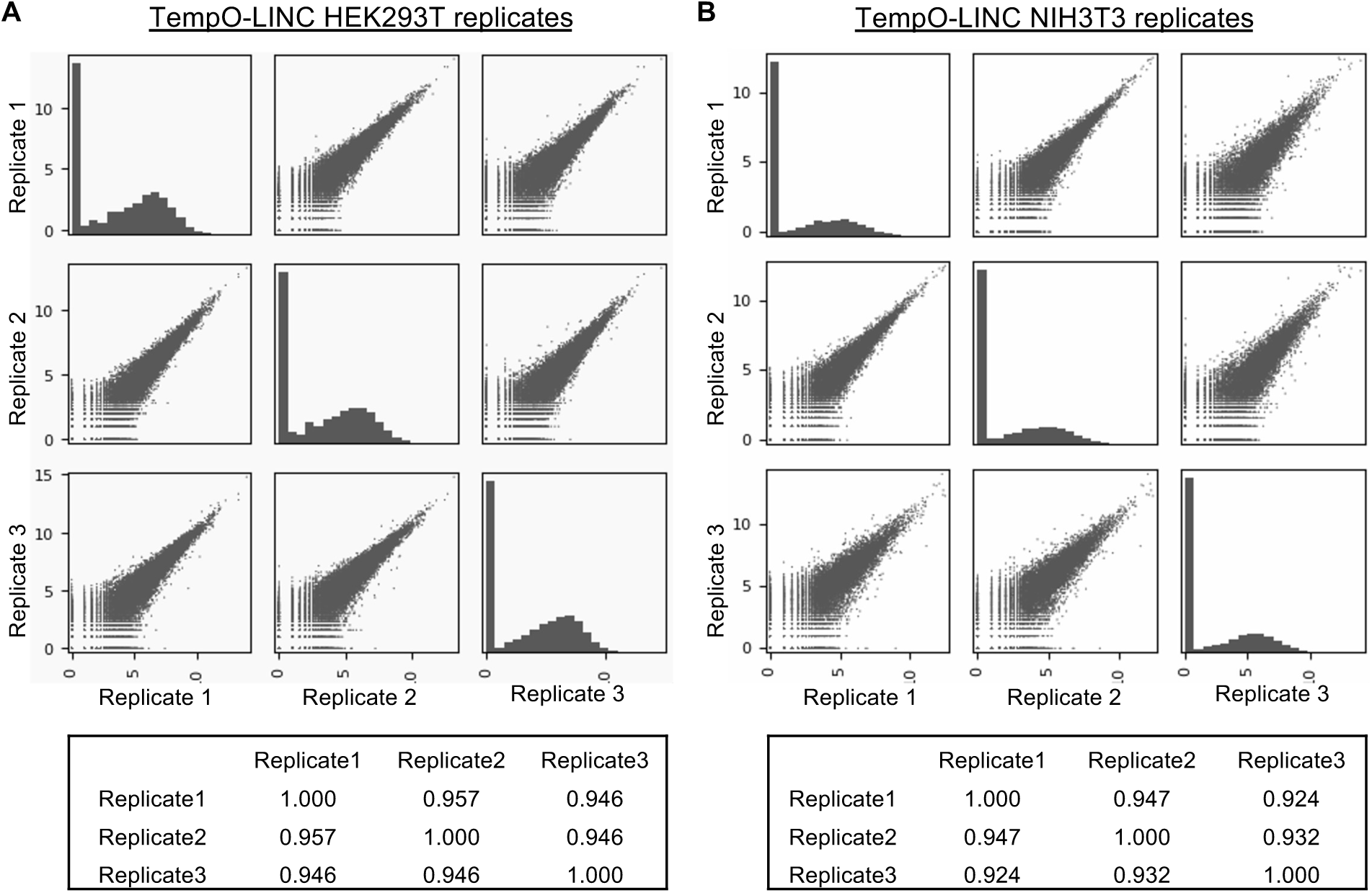
(A) The scatterplot matrix shows the correlation between the sum of the log2 transformed read counts from 50 random human HEK293T cells across three TempO-LINC replicates. Plots reported in the diagonal of the matrix show the distribution of the log2 transformed read counts for each replicate (x axis = log2 transformed sum of counts from 50 random cells, y axis: number of genes). Pearson *r* values are reported in the table below the scatterplot matrix. (B) Identical scatterplot matrix shown for the sum of log2 transformed read counts from 50 random mouse NIH3T3 cells in the three replicates.

**Supplemental Figure 3.**
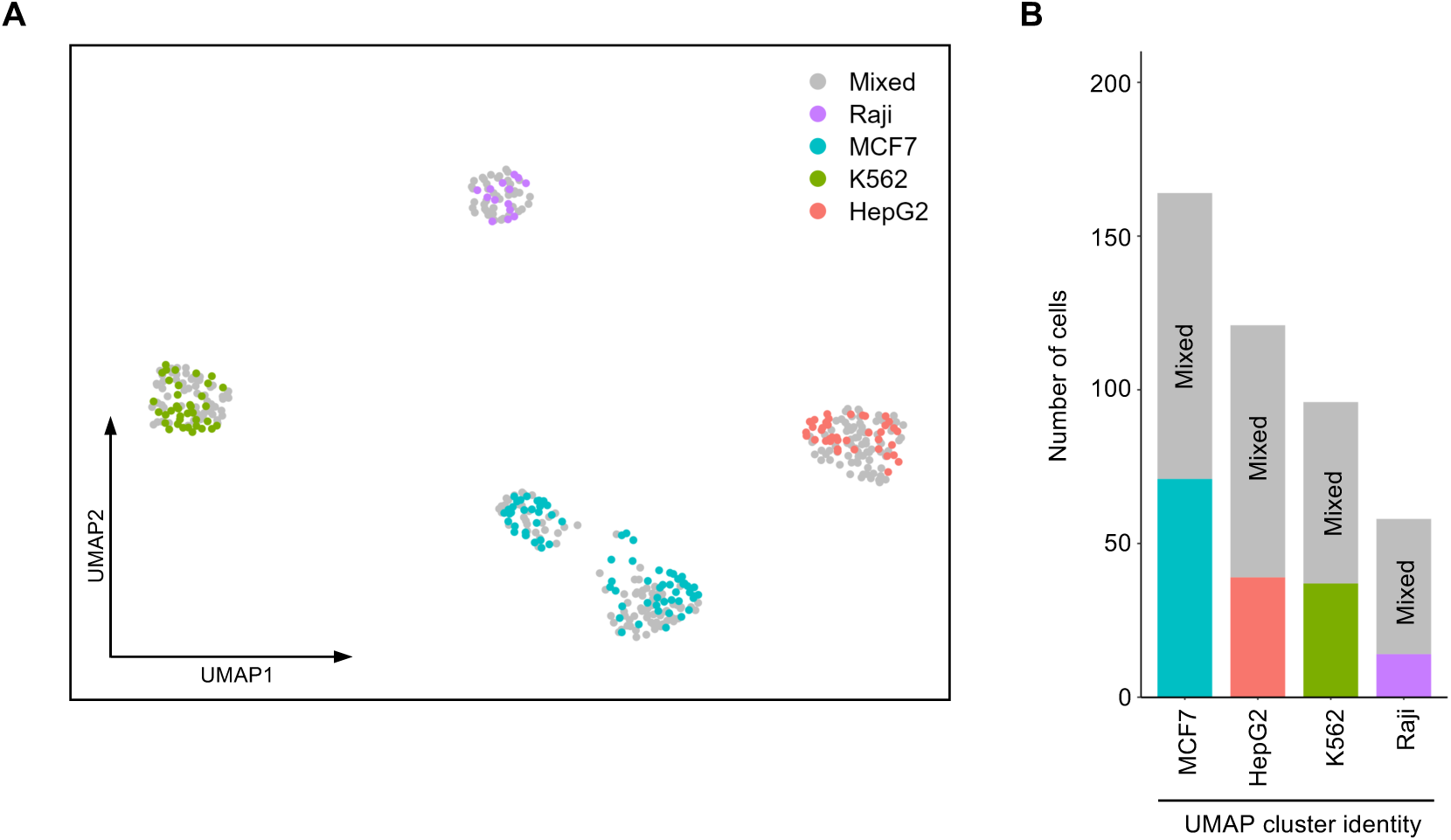
(A) Human MCF7, Raji, K562 and HepG2 cell lines as well as a mixture of all four cell lines (Mixed) were simultaneously barcoded with TempO-LINC for single-cell transcriptome profiling. 439 total cells were identified with a UMAP plot showing four distinct populations of cells identified following unsupervised clustering. Based on the Barcode 1 identity, cells for mixed and independent cell line samples were color coded on the UMAP, demonstrating that TempO-LINC can correctly resolve cell type identity from mixed populations of human cells. (B) Bar plot showing the number of cells associated with each of the 4 major UMAP populations. The number of cells associated with “Mixed” (gray fraction) or cell line-specific barcode sample identities can be seen for each bar on the plot. The precise cell numbers were: 278 total Mixed cells identified and found distributed into all 4 of the cell cluster populations, 71 MCF7 cells, 39 HepG2 cells, 14 Raji cells and 37 K562 cells.

**Supplemental Figure 4.**
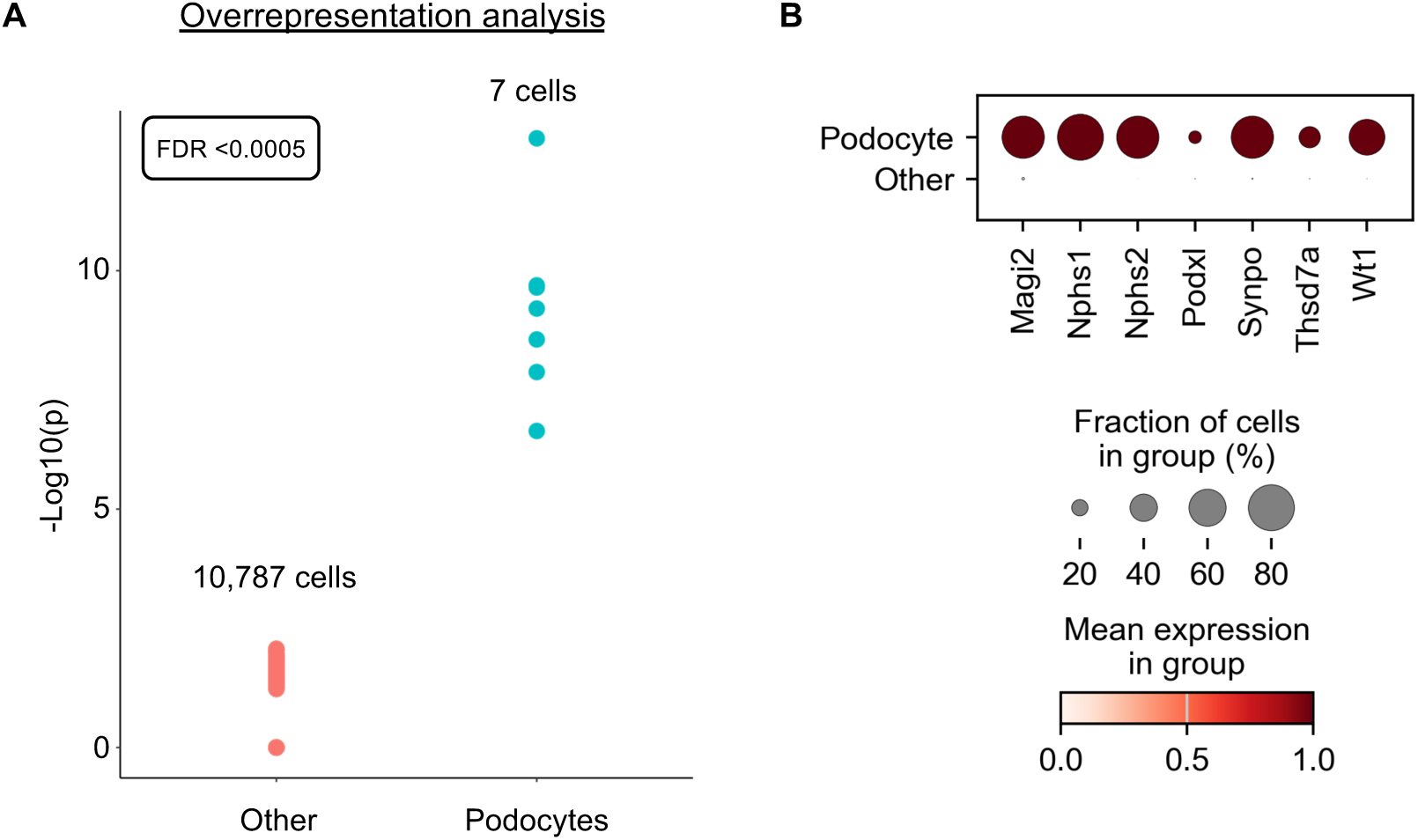
Identification of Podocytes from Kidney Tissue. (A) Identification of podocytes within the mouse kidney was performed using overrepresentation analysis utilizing a seven podocyte marker gene set consisting of Nphs1, Nphs2, Podxl, Wt1, Magi2, Synpo and Thsd7a. The false discovery rate (FDR) value was calculated using the Benjamini & Hochberg adjustment method. (B) Dot plot analysis showing expression of seven known molecular markers for podocytes across either the identified podocytes or the 10,787 “other” cells resulting from the overrepresentation analysis in panel “A”.

**Supplemental Figure 5.**
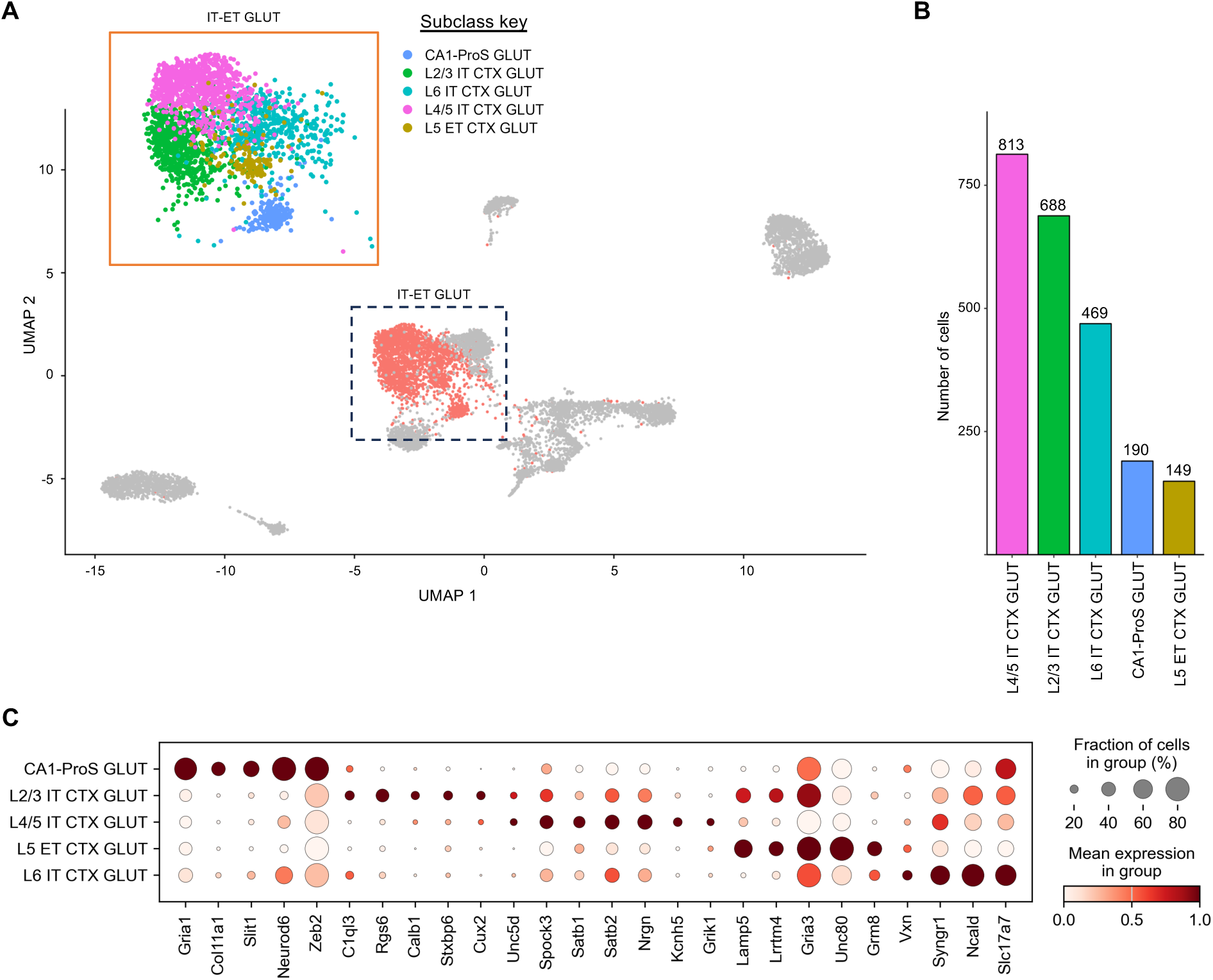
Neuronal Subclass Identification within the IT-ET GLUT Cluster. (A) Increasing the cluster resolution on the Seurat generated Figure 4 UMAP demonstrates that 5 additional neuronal cell subclusters can be identified within the IT-ET GLUT population. Dashed box indicates the region on the UMAP in Figure 4 that is magnified in the orange box. IT-ET subclasses identified and annotated include: L2/3 IT CTX GLUT, L4/5 IT CTX GLUT, L6 IT CTX GLUT, L5 ET CTX GLUT and CA1-ProS GLUT. (B) Cell numbers for each subclass are plotted. (C) Dot plot showing expression of 26 subclass specific biomarkers within the 5 identified IT-ET GLUT subclusters.

